# Learning quantitative sequence-function relationships from high-throughput biological data

**DOI:** 10.1101/020172

**Authors:** Gurinder S. Atwal, Justin B. Kinney

**Affiliations:** Simons Center for Quantitative Biology, Cold Spring Harbor Laboratory; Simons Center for Quantitative Biology, Cold Spring Harbor Laboratory Cold Spring Harbor, NY 11724

## Abstract

Understanding the transcriptional regulatory code, as well as other types of information encoded within biomolecular sequences, will require learning biophysical models of sequence-function relationships from high-throughput data. Controlling and characterizing the noise in such experiments, however, is notoriously difficult. The unpredictability of such noise creates problems for standard likelihood-based methods in statistical learning, which require that the quantitative form of experimental noise be known precisely. However, when this unpredictability is properly accounted for, important theoretical aspects of statistical learning which remain hidden in standard treatments are revealed. Specifically, one finds a close relationship between the standard inference method, based on likelihood, and an alternative inference method based on mutual information. Here we review and extend this relationship. We also describe its implications for learning sequence-function relationships from real biological data. Finally, we detail an idealized experiment in which these results can be demonstrated analytically.

## 1 Introduction

A major long-term goal in biology is to understand how biological function is encoded within the sequences of DNA, RNA, and protein. The canonical success story in this effort is the genetic code. Given an arbitrary sequence of messenger RNA, the genetic code allows us to predict with near certainty what peptide sequence will result.

But there are many other biological codes we would like to learn. How does the DNA sequence of a promoter or enhancer encode transcriptional regulatory programs? How does the sequence of pre-mRNA govern which exons are kept and which are removed from the final spliced mRNA? How does the peptide sequence of an antibody govern how strongly it binds to target antigens? For each of these questions, we aim to learn a sequence-function relationship: given the sequence of a biological heteropolymer, what will that molecule end up actually doing?

A major difference between the genetic code and these other codes is that while the former is qualitative in nature, the latter govern biological processes that are inherently quantitative. For example, the rate of transcription from a promoter can range continuously over orders of magnitude, and the value of this rate can be finely tuned by adjusting the DNA sequence of the promoter. Learning these biological codes thus requires theoretical and computational methods to infer quantitative sequence-function relationships from data.

Experimental methods for measuring sequence-function relationships have improved dramatically over the last 15 years. The availability of microarray-based experiments [1, 2], for example, made it possible to accurately measure the sequence-specificity of individual DNA-binding proteins, such as transcription factors. More recently, multiple techniques (e.g., [3–5]) for simultaneously measuring the activity of many different transcriptional regulatory sequences have been developed. Below we discuss one of these methods, Sort-Seq [5], in greater detail.

These experimental methods, which can measure the activities of many different biological sequences at once, are very unlike conventional experiments in physics. Such high-throughput biological experiments are typically very noisy and rarely provide direct readouts for the quantities that one cares about. Moreover, these measurements generally exhibit substantial day-to-day variability. However, as shown in this paper, it is still possible to precisely learn quantitative sequence-function relationships from such data, even though the noise characteristics of these data are difficult or impossible to characterize up front.

The ability to do this reflects subtle but important distinctions between two objective functions used for statistical inference: (i) likelihood, which requires *a priori* knowledge of the experimental noise function and (ii) mutual information, a fundamental quantity in information theory [6] that does not require a noise function. In contrast to the conventional wisdom that more experimental measurements will improve the model inference task, we show that there are instances in which the popular maximum likelihood approach will never learn the right model, even in the infinite data limit. Model inference based on mutual information does not suffer from this ailment, but it is unable to determine the values of a small subset of model parameters known as “diffeomorphic modes” [7]. This inability of mutual information to pin down parameters along diffeomorphic modes does not indicate a problem with mutual information, but rather reflects a fundamental distinction between how diffeomorphic and nondiffeomorphic model parameters are constrained by data.

In this paper we detail these two distinct inference approaches in the context of learning sequence-function relationships and elaborate their relationship to one another. We also work through an explicit analytical example of inference in an idealized Sort-Seq experiment. This example highlights the differences between likelihood- and mutual-information-based approaches to inference, as well as the emergence of diffeomorphic modes.

It should be noted that the inference of receptive fields in sensory neuroscience is another area of biology in which mutual information has proved useful as an objective function, and that work in this area has also provided important insights into basic aspects of machine learning [8–12]. This body of work, however, has largely avoided in-depth discussions of how mutual information relates to likelihood, which is a major focus of the present paper.

## 2 Sort-Seq and other experimental methods

We begin by describing Sort-Seq [5], one of the recently developed experimental techniques for measuring quantitative sequence-function relationships. This assay uses flow cytometry and high-throughput DNA sequencing to simultaneously measure the activity of a large number of different transcriptional regulatory sequences. It was first used for characterizing the biophysical basis of transcriptional regulation in *E. coli*. This basic approach has since been adapted for studying multiple aspects of gene regulation in bacteria *E. coli* [13], yeast [14], and human cells [15]. Sort-Seq-like methods have also been used to study protein sequence-function relationships in various systems [16, 17].

First, a DNA plasmid is created in which a regulatory sequence of interest is placed upstream of a fluorescent reporter gene, such as green fluorescent protein (GFP). A large library of plasmids is then created by replacing this regulatory sequence with variant regulatory sequences containing randomly scattered substitution mutations. The resulting plasmids are then inserted into cells, which are allowed to grow under expression-inducing conditions. Cells then undergo fluorescence-activated cell sorting (FACS). This procedure sorts the cells into a small number of “batches” based on each cell’s measured fluorescence. FACS machines can sort thousands of cells a second, so it is not difficult to sort *∼*10^6^ cells into each batch.

Finally, the mutated regulatory sequences within each batch are amplified using PCR and then sequenced. The resulting data set consists of a large number of mutant regulatory sequences *S*, each with a corresponding measurement *M* (the batch in which the sequence is found). In the original experiment [5], *∼*2.5 *×* 10^5^ independent measurements were taken. Advances in DNA sequencing have since made it possible to accumulate much more data, and it is no longer difficult to assay the activities of millions of sequences in a single experiment.

Fig. 2 depicts biophysical models of transcriptional regulation that were learned from Sort-Seq data in the study of [5]. These illustrate just a few of the quantitative sequence-function relationships that Sort-Seq and related experiments can elucidate. In the Sort-Seq experiments of [5], a 75 bp region of the *E. coli lac* promoter was interrogated. This region contains binding sites for two regulatory proteins, the RNA polymerase holoenzyme (RNAP) and the cAMP receptor protein (CRP) transcription factor (Fig. 2a). The resulting Sort-Seq data was used to learn “energy matrix” models for the sequence-dependent binding energies of CRP and RNAP (Fig. 2b,c). These data were also used to infer a biophysical model describing how these two proteins interact (Fig. 2d). By fitting these quantitative sequence-function relationships to data, it was possible to learn physically meaningful values for the *in vivo* molecular interaction energies that form the mechanistic basis for transcriptional regulation in this system. The specific way such models were fit to data is discussed in more detail in section 6.3 of this paper.

**Fig. 1.**
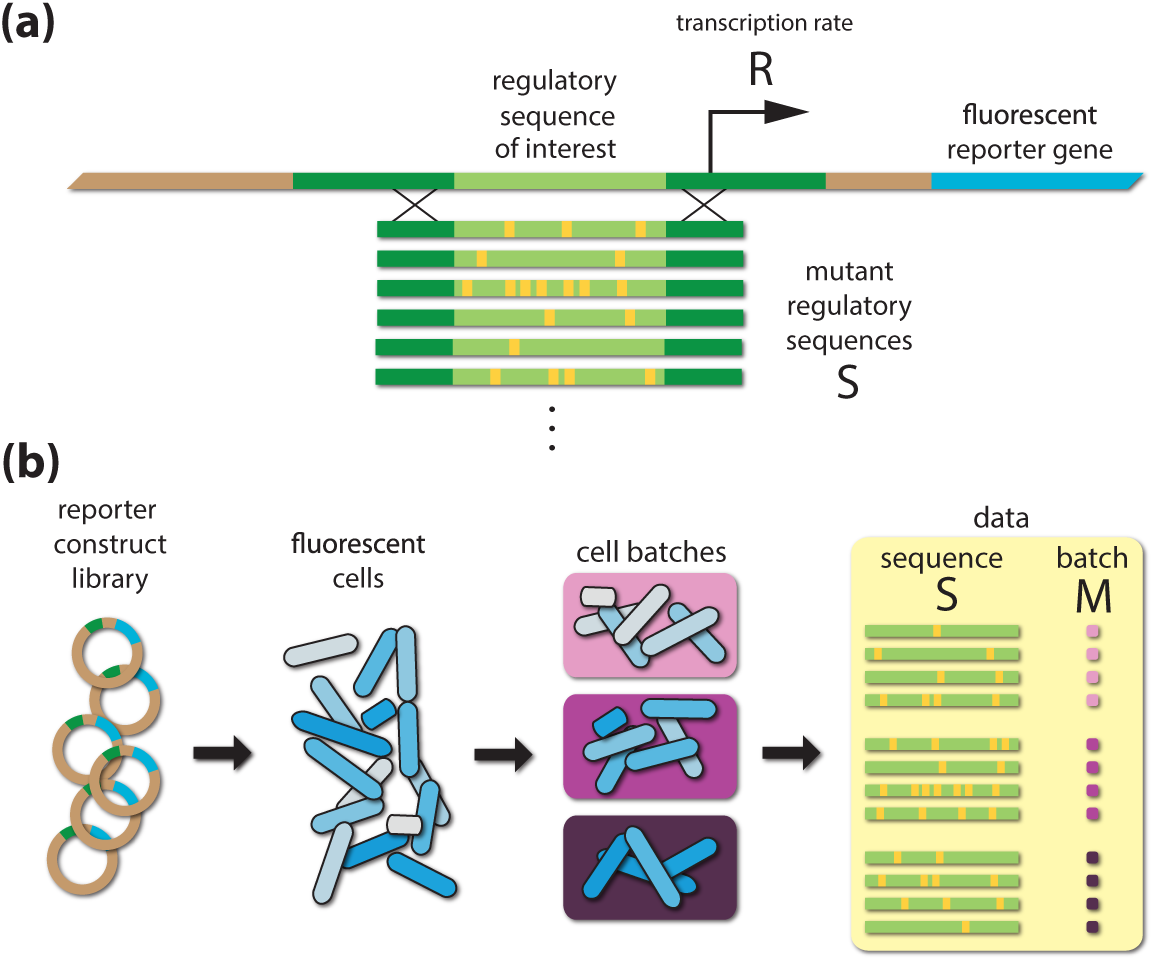
Overview of the Sort-Seq assay [5]. (a) A regulatory sequence of interest is inserted into a plasmid upstream of a fluorescent reporter gene. This regulatory sequence is then replaced with variant sequences, each having multiple scattered substitution mutations. (b) The resulting plasmid library is put into cells. Cells are cultured under inducing conditions, then sorted into batches according to fluorescence. The variant regulatory sequences in each batch of sorted cells are then sequenced *en masse*. The resulting data set is comprised of a large number of variant sequences *S*, each assigned to a batch *M*. Figure adapted from [5].

**Fig. 2.**
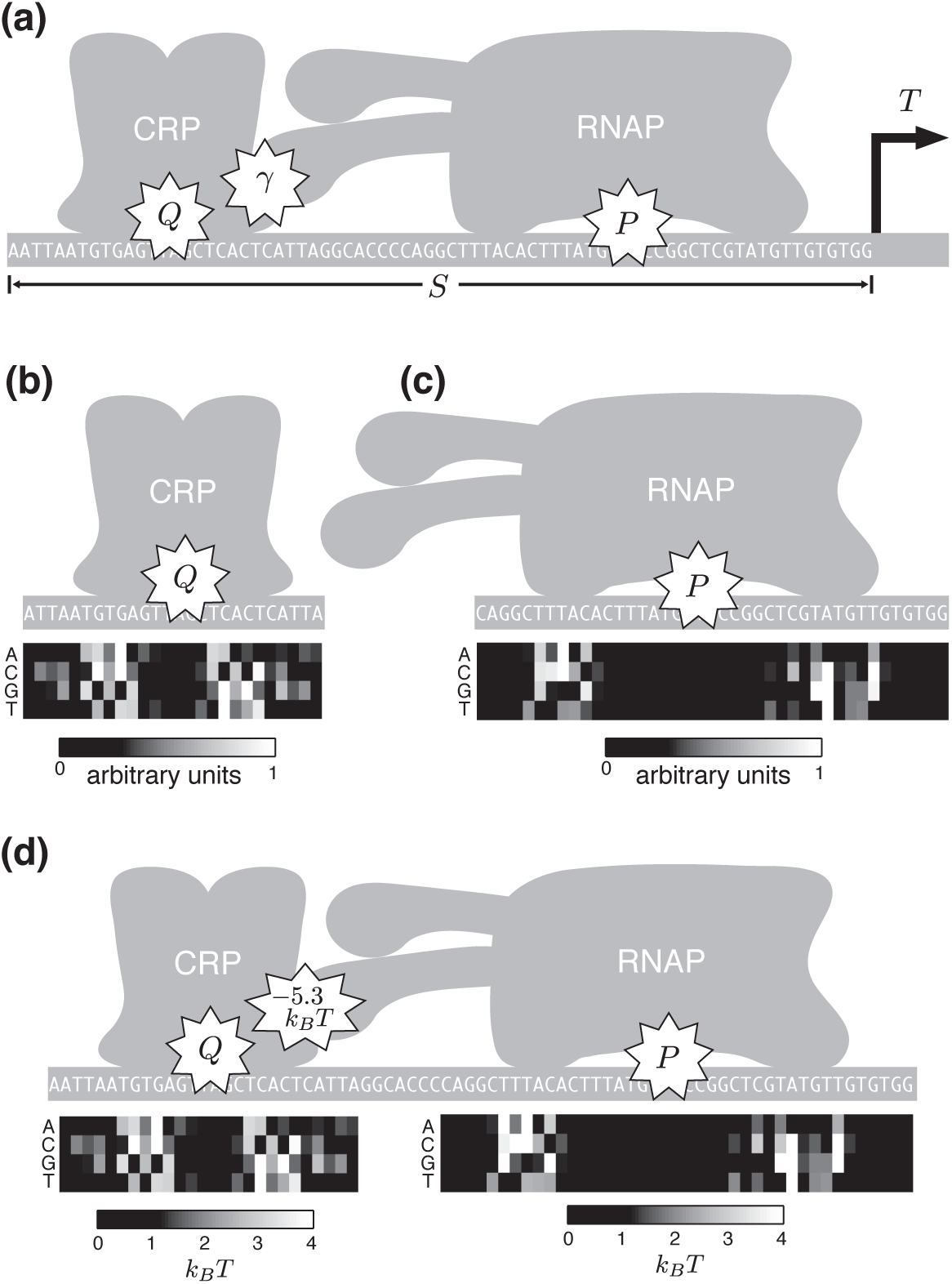
Sequence-function relationships fit to Sort-Seq data in [5]. (a) The authors inferred a biophysical model of transcriptional regulation by the *lac* promoter. This biophysical model included two “energy matrix” models, one describing the sequence-dependent binding energy of CRP (*Q*), and one describing the binding energy of RNAP (*P*). It also included a value for the interaction energy *γ* between these two proteins. (b) Fitting only the CRP energy matrix resulted in an energy matrix with unknown scale. (c) Similarly, fitting only the RNAP energy matrix resulted in an unknown energy scale. (d) Fitting the full biophysical model recovered both CRP and RNAP energy matrices in meaningful energy units, as well as a value for the protein-protein interaction energy *γ*. Figure adapted from [7].

Before proceeding, we briefly describe other experimental methods that can produce data needing similar analysis. One alternative to sorting sequences is to directly quantify the abundance of transcripts produced from different regulatory sequences using RNA-Seq. Ways of using this approach to dissect regulatory sequences *in vitro* [4] and *in vivo* [18–20] have been described. Another method is to perform a selection assay that enriches for sequences according to some measured activity, then to sequence both the selected and non-selected batches of sequences [3, 21, 22].

## 3 Inference using likelihood

The inference of quantitative sequence-function relationships from Sort-Seq data can be phrased as follows. We have a data set consisting of a large number of sequences 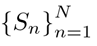, each sequence *S* having a corresponding measurement *M*. This process is stochastic: due to experimental noise, different measurements of the same sequence *S* can yield different values for *M*. Our experiment therefore has the following form

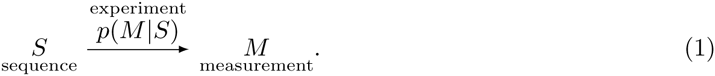

If we have an explicit parametric form for *p*(*M |S*), we can learn the values of the parameters by maximizing the per-datum log likelihood,

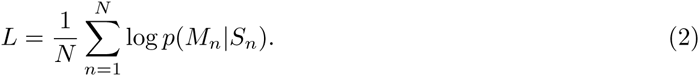

Here we have assumed that the measurements for each sequence are independent. In regression problems such as this, one introduces an additional layer of structure. In this case, we expect the measurement *M* of each sequence *S* to be a noisy readout of some underlying activity *R* that is a deterministic function of that sequence; we call this function our “model”, and denote it *θ*(*S*). This model is ultimately what we care about. The noisiness of the experiment is then characterized by a “noise function” *π*(*M* |*R*).Our experiment is thus represented by

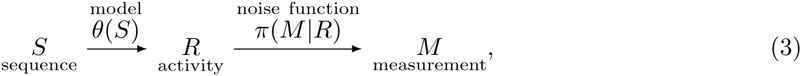

where *p*(*M*|*S*) = *π*(*M*∣*θ*(*S*)). The corresponding likelihood is

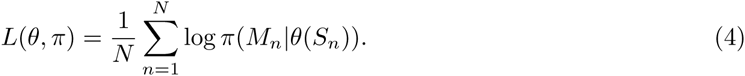

The model we adopt for our experiment therefore has two components: *θ*, which we want to learn from data, and *π*, which we do not really care about.

Standard statistical regression requires that the noise function *π* be specified up-front. *π* can be learned either by performing separate calibration experiments, or by assuming a functional form based on an educated guess. This can be problematic, however. Consider inference in the large data limit, (*N* → ∞) which is illustrated in Fig. 3. Likelihood is determined by both the model *θ* and the noise function *π* (Fig. 3a). If we know the correct noise function *π^*^* exactly, then maximizing *L*(*θ, π^*^*) over *θ* is guaranteed to recover the correct model *θ^*^*. However, if we assume an incorrect noise function *π^′^*, maximizing likelihood will typically recover an incorrect model *θ^′^* (Fig. 3b).

**Fig. 3.**
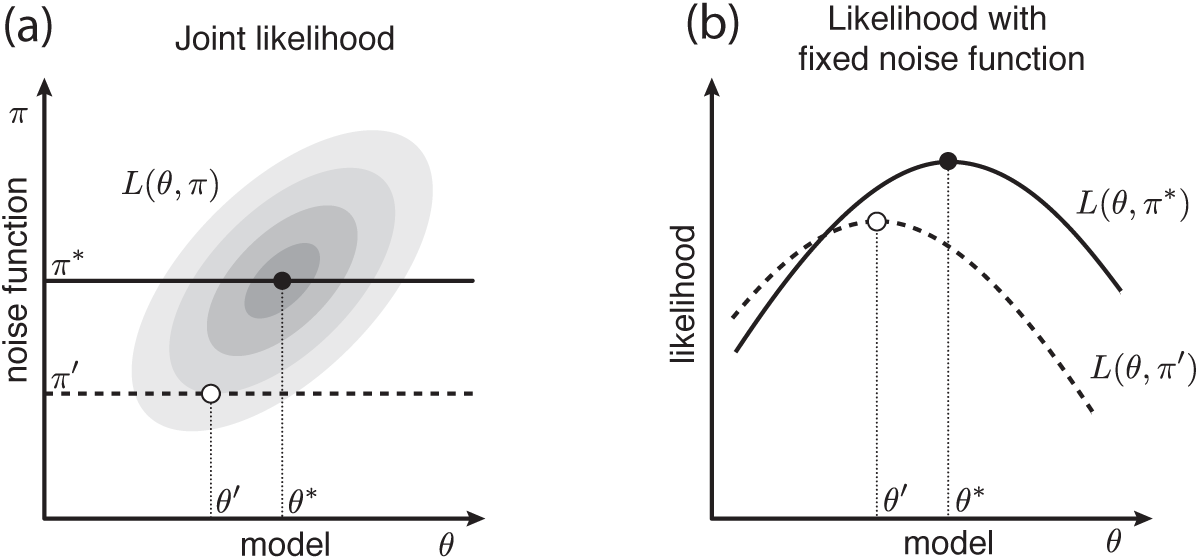
Schematic illustration of how likelihood *L*(*θ, π*) depends on the model *θ* and the noise function *π* in the *N→ ∞* limit. (a) *L* will typically have a correlated dependence on *θ* and *π*. (b) If *π* is set equal to the correct noise function *π^*^*, then *L* will be maximized by the correct model *θ^*^*. However, if *π* is set to an incorrect noise function *π^′^*, *L* will typically attain a maximum at an incorrect model *θ^′^*. Figure adapted from [7].

## 4 Inference using mutual information

Information theory provides an alternative inference approach. Suppose we hypothesize a specific model *θ*, which gives predictions *R*. Furthermore, denote the true model *θ^*^* and the corresponding true activity *R*^*\**^. The dependence between *S*, *M*, *R*^*\**^, and *R* will then follow the graphical model,

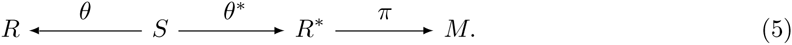

From the simple fact that *M* depends on *R* through the value of *R*^*\**^ any dependence measure 𝒟 that satisfies the Data Processing Inequality (DPI) must satisfy

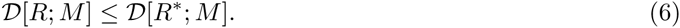

Therefore, in the set of possible models *θ*, the true model is guaranteed to globally maximize the objective function 𝒟 (θ) ≡ 𝒟 [R;M].

One particularly relevant dependence measure satisfying DPI is mutual information. Mutual information plays a fundamental role in information theory [6]. It also provides a self-equitable measure of statistical association [23]. In the case of Sort-Seq experiments, which have continuous *R* and discrete *M*, mutual information is given by

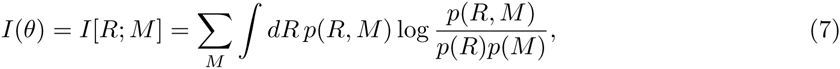

where *p*(*M, R*) is the distribution of activity predictions and measurements resulting from the model *θ*. Note, in particular, that computing mutual information does not depend on the assumption of any noise function *π*. Rather, if one is able to estimate *p*(*M, R*) from a finite sample of data, mutual information can be used as an objective function for determining *θ* without assuming any noise function *π*.

It should be noted that there are multiple dependence measures 𝒟 that satisfy DPI. One might wonder whether maximizing multiple different dependence measures would improve on the optimization of mutual information alone. The answer is not so simple. In [7] it was shown that if the correct model *θ^*^* is within the space of models under consideration, then, in the large data limit, maximizing mutual information is equivalent to simultaneously maximizing every dependence measure that satisfies DPI. On the other hand, one rarely has any assurance that the correct model *θ^*^* is within the space of parameterized models one is considering. In this case, considering different DPI-satisfying measures might provide a test for whether *θ^*^* is noticeably outside the space of parameterized models. To our knowledge, this potential approach to the model selection problem has yet to be demonstrated.

## 5 Relationship between likelihood and mutual information

An alternative inference approach is to admit that we do not know the noise function *π a priori*, and to fit *both θ* and *π* simultaneously by maximizing *L*(*θ, π*) over this pair. It is easy to see why this makes sense: the division of the inference problem into first measuring *π*, then learning *θ* using that inferred *π*, is somewhat artificial. The process that maps *S* to *M* is determined by both *θ* and *π* and thus, from a probabilistic point of view, it makes sense to maximize likelihood over both of these quantities simultaneously.

We now show that, in the large *N* limit, maximizing likelihood over both *θ* and *π* is equivalent to maximizing the mutual information between model predictions and measurements. Here we follow the argument given in [7]. In the large *N* limit, likelihood can be decomposed into three terms

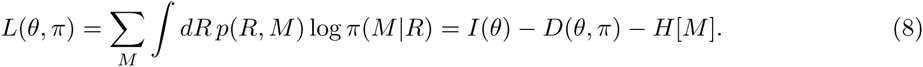

where

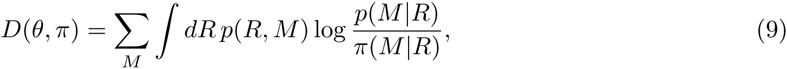

is the Kullback-Leibler divergence between the assumed noise function *π* and the observed noise function *p*(*M|R*), and *p*(*M|R*), and *H*[*M*] = − ∑_*M*_ *p*(*M*) log *p*(*M*) is the entropy of the measurements, which does not depend on *θ*. To maximize *L*(*θ, π*), it therefore suffices to maximize *I*(*θ*) over *θ* alone, then to set the noise function *π*(*M |R*) equal to the empirical noise function *π*(*M*|*R*) which causes *D*(*θ, π*) to vanish.

Thus, when we are uncertain about the noise function *π*, we need not despair. We can, if we like, simply learn *π* at the same time that we learn *θ*. We need not explicitly model *π* in order to do this; it suffices instead to maximize the mutual information *I*(*θ*) over *θ* alone.

The connection between mutual information and likelihood can further be seen in a quantity called the “noise-averaged” likelihood. This quantity was first described for the analysis of microarray data [24]; see also [7]. The central idea is to put an explicit prior on the space of possible noise functions, then compute likelihood after marginalizing over these noise functions. Explicitly, the per-datum log noise-averaged likelihood *L*_*na*_*(θ)* is related to *L(θ,π)* via

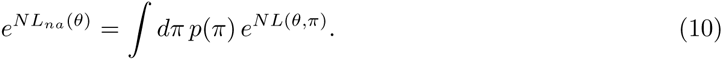

Under fairly general conditions, one finds that noise-averaged likelihood is related to mutual information via

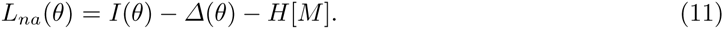

Here, *Δ*(*θ*) is a quantity that vanishes in the *N* limit, and thus becomes irrelevant for the inference problem on sufficiently large data sets.

## 6 Diffeomorphic modes

Mutual information has a mathematical property that is important to account for when using it as an objective function: the mutual information between any two variables is unchanged by an invertible transformation of either variable. So if a change in model parameters, *θ → θ^1^*, results in changes in model predictions *R → R^′^* that preserves the rank order of these predictions, then

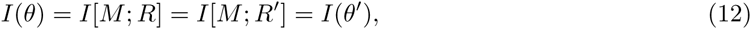

and *θ* and *θ*^′^ are judged to be equally valid.

By using mutual information as an objective function, we are therefore unable to constrain any parameters of *θ* that, if changed, produce invertible transformations of model predictions. Such parameters are called “diffeomorphic parameters” or “diffeomorphic modes” [7]. The distinction between diffeomorphic parameters and nondiffeomorphic parameters is illustrated in Fig. 4.

**Fig. 4.**
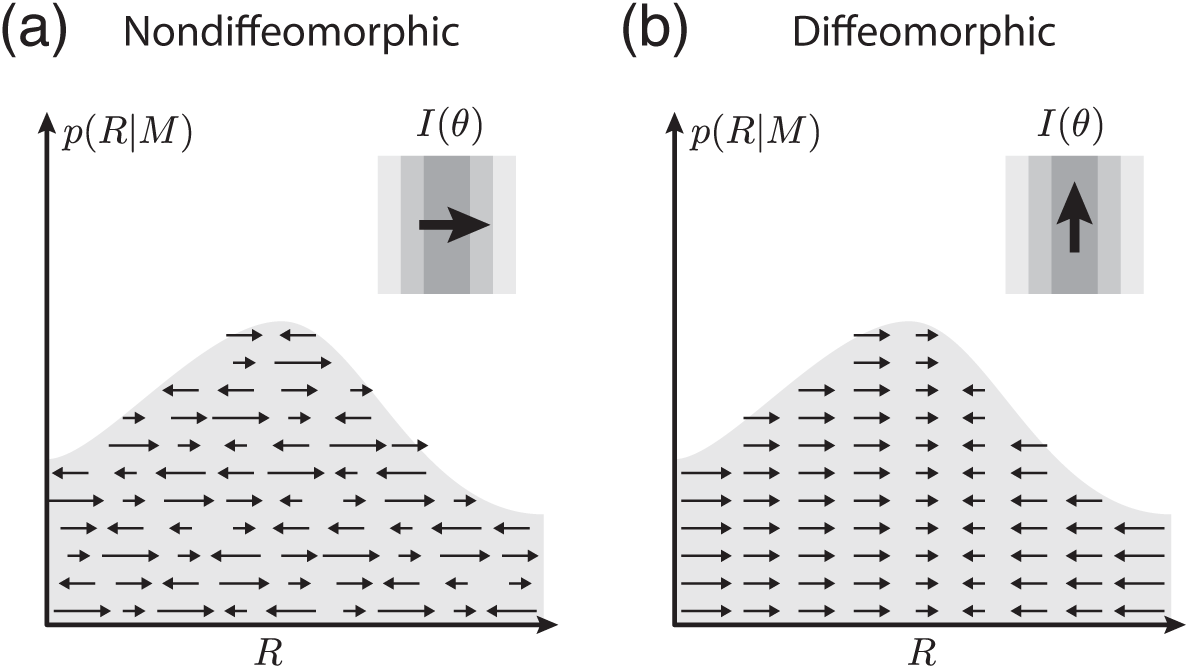
Illustration of diffeomorphic and nondiffeomorphic parameters. (a) Changing the value of a nondiffeomorphic parameter of *θ* results in a sort of “diffusion” in which the *R* values assigned to different sequences change rank order. Such changes will typically alter the mutual information *I*(*θ*) = *I*[*R*; *M*]. (b) Changing the value of a diffeomorphic parameter, however, results in a “flow” of *R* values that maintains their rank order. Such flows in *R*-space leave *I*(*θ*) unchanged.

### 6.1 Criterion for diffeomorphic modes

Following [7], we now derive a criterion that can be used to identify all of the diffeomorphic modes of a model *θ*.^1^ Consider an infinitesimal change in model parameters *θ → θ + dθ*, where the components of *dθ* are specified by

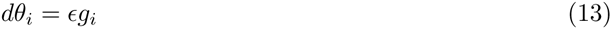

for some vector *g*_*i*_ in *θ*-space. This change in *θ* will produce a corresponding change in model predictions *R → R* + *dR*, where

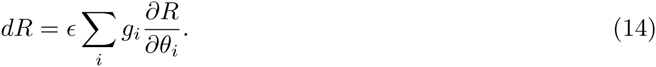

In general, the derivative *∂R/∂θ_i_* can have arbitrary dependence on the underlying sequence *S*. However, this transformation will preserve the rank order of *R*-values if and only if *dR* is the same for all sequences having the same value of *R*. The change *dR* must therefore be a function of *R*, and have no other dependence on *S*. A diffeomorphic mode is a vector field *g*(*θ*) that has this property at all points in parameter space. Specifically, a vector field *g*(*θ*) is a diffeomorphic mode if and only if there is a function *h*(*R, θ*) such that

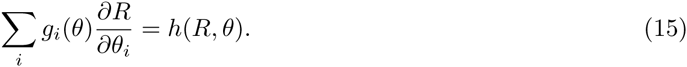

### 6.2 Diffeomorphic modes of linear models

As a simple example, consider a situation in which each sequence *S* is a *D*-dimensional vector, and *R* is an affine function of *S*, i.e.

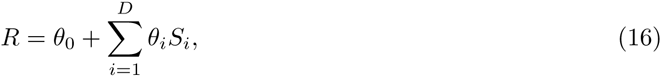

for model parameters *θ* = {*θ*_0_, *θ*_1_, … *θ*_*D*_} The criterion (Eq. (15)) for a vector field *g* being a diffeomorphic mode then gives

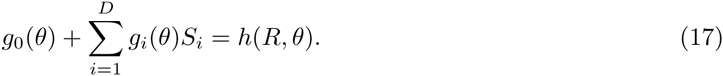

Because the left hand side is linear in *S*, and *R* is linear in *S*, the function *h*(*R, θ*) must be linear in *R*. Thus, *h* must have the form

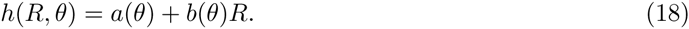

So at each *θ*, the vector field *g*(*θ*) has at most two degrees of freedom, which correspond to the values of *a*(*θ*) and *b*(*θ*). Specifically,

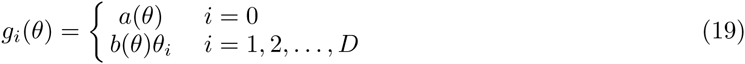

The *a* component of *g* corresponds to adding a constant to *R*, while the *b* component corresponds to multiplying *R* by a constant.

Note that if we had instead chosen 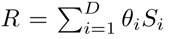 i.e. left out the constant component *θ*_0_, then there would be only one diffeomorphic mode, corresponding to multiplication of *R* by a constant. This fact will be used when we analyze the Gaussian selection model in Section 8.

### 6.3 Diffeomorphic modes of a biophysical model of transcriptional regulation

Diffeomorphic modes can become less trivial in more complicated situations. Consider the biophysical model of transcriptional regulation by the *E. coli lac* promoter (Fig. 2). This model was fit to Sort-Seq data in [5]. The form of this model is as follows. Let *S* denote a 4 × *D* matrix representing a DNA sequence of length *D* and having elements

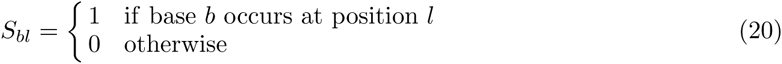

where *b* ϵ {*A;C; G; T*} and *l* = 1, 2, *… D*. The binding energy *Q* of CRP to DNA was modeled in [5] as an “energy matrix”: each position in the DNA sequence was assumed to contribute additively to the overall energy. Specifically,

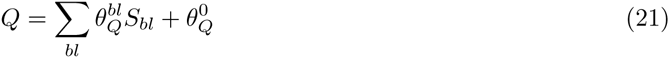

where 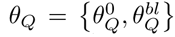 are the parameters of this energy matrix. Similarly, the binding energy P of RNAP to DNA was modeled as an energy matrix

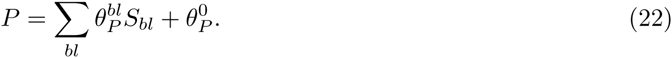

Both energies were taken to be in thermal units (*k*_*B*_*T*). The rate of transcription *T* resulting from these binding energies was taken to be proportional to the occupancy of RNAP at its binding site. This is given by

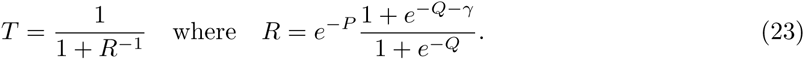

where **γ** is the interaction energy between CRP and RNAP (again in units of *k*_*B*_*T*).

Because the binding sites for CRP and RNAP do not overlap, one can learn the parameters *θ_Q_* and *θ_P_* from data separately by independently maximizing *I*[*Q*; *M*] and *I*[*P* ; *M* ]. The results of doing this are shown in Fig. 2b,c. Note, however, that that the overall scale of each energy matrix remains undetermined, as does each chemical potential 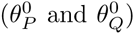. The reason is that the energy scale and chemical potential are diffeomorphic modes of energy matrix models and therefore cannot be inferred by maximizing mutual information.

However, if *Q* and *P* are inferred together by maximizing *I*[*T* ; *M*] instead, one is now able to learn both energy matrices with a physically meaningful energy scale. The chemical potential of CRP, 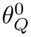, is also determined. The only parameter left unspecified is the chemical potential of RNA polymerase, 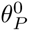. The reason for this is that, while *Q* and *P* together have four diffeomorphic modes, *T* only has one. In the formula for *T*, the energies *P* and *Q* combine in a nonlinear way. This nonlinearity eliminates three of the four diffeomorphic modes. See [7] for the derivation of this result.

### 6.4 Conjugate noise modes

Diffeomorphic modes can be thought of as being “conjugate” to certain modes of the noise function. Consider a diffeomorphic transformation *θ*→*θ*′ Because this transformation is diffeomorphic, the predictions *R*^*′*^ of *θ^′^* will be related to the predictions *R* of *θ* by some invertible function *f* :

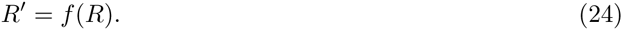

Now suppose that, at the same time we perform the transformation *θ → θ^′^*, we also transform our noise function *π → π^′^* where *π^′^* is defined by

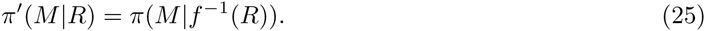

The resulting transformation (*θ, π*) *→* (*θ^1^, π^′^*) leaves *p*(*M |S*) invariant, since

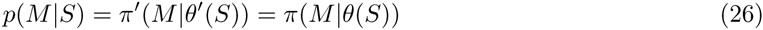

for all sequences *S* and measurement values *M*. In what follows, we refer to the noise function transformation *π* → *π*′ as being conjugate to the model transformation *θ*→ *θ*′

It is useful to think of the diffeomorphic parameters of *θ* and the conjugate parameters of *π* in terms of the Venn diagram shown in Fig. 5. Typically, changes to most of the parameters of *θ* will produce transformations of *p*(*M | S*) that cannot be achieved by any possible changes to *π*. The reverse is also true. However, changes to any diffeomorphic parameter of *θ* will produce the same transformations of *p*(*M | S*) as a corresponding change to the conjugate parameter of *π*. Thus, if we think of *θ* and *π* in terms of the transformations of *p*(*M | S*) that they generate, then the diffeomorphic parameters of *θ* and the conjugate parameters of *π* are seen to lie within the intersection of *θ* and *π*.

**Fig. 5.**
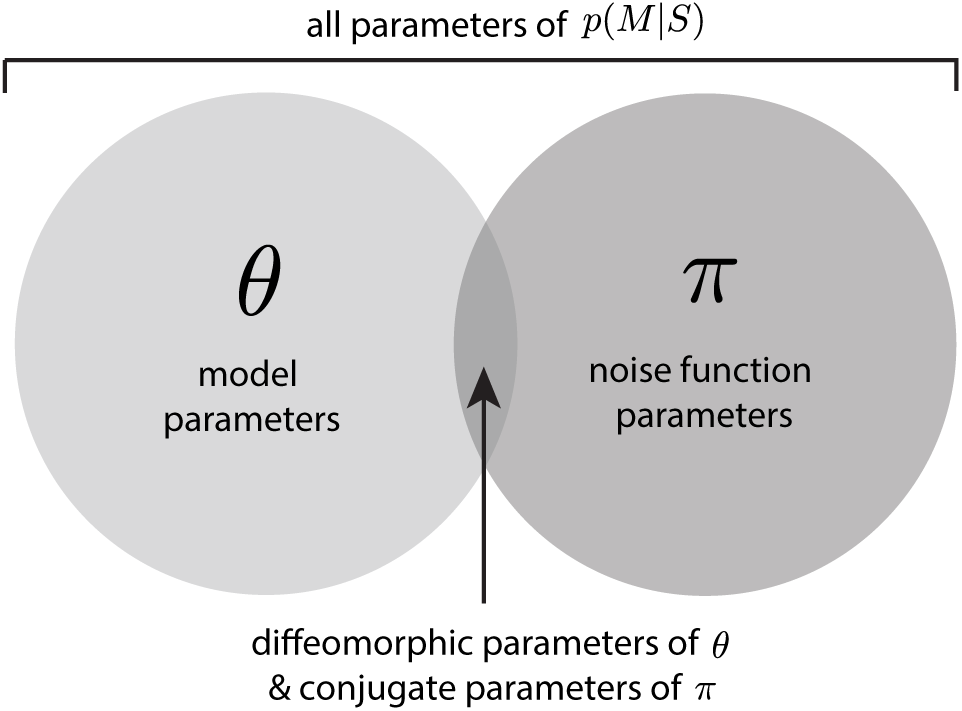
Venn diagram illustration of which parameters control which degrees of freedom of *p*(*M| S*). Typically, altering the model *θ* will change *p*(*M |S*) in a way that cannot be be achieved by altering the noise function *π*. However, changes to the diffeomorphic parameters in *θ* will change *p*(*M S*) in a manner identical to changing the conjugate parameters of *π*. Thus, it is useful to think of diffeomorphic model parameters and their conjugate noise function parameters as lying within the intersection of *θ* and *π*.

This finding has an intuitive interpretation: there is ambiguity in how we divide our experiment up into an activity model *θ* and a noise function *π*. This ambiguity is parameterized by the diffeomorphic modes of *θ* and their conjugate parameters of *π*. In what follows, we will see that being cognizant of this ambiguity is critical for correctly inferring activity models from data.

## 7 Inference using likelihood, noise-averaged likelihood, and mutual information

The diffeomorphic modes of parametric models respond to data in different ways than do nondiffeomorphic modes. This is illustrated in Fig. 6. Consider the range of parameter values consistent with the posterior distribution

**Fig. 6.**
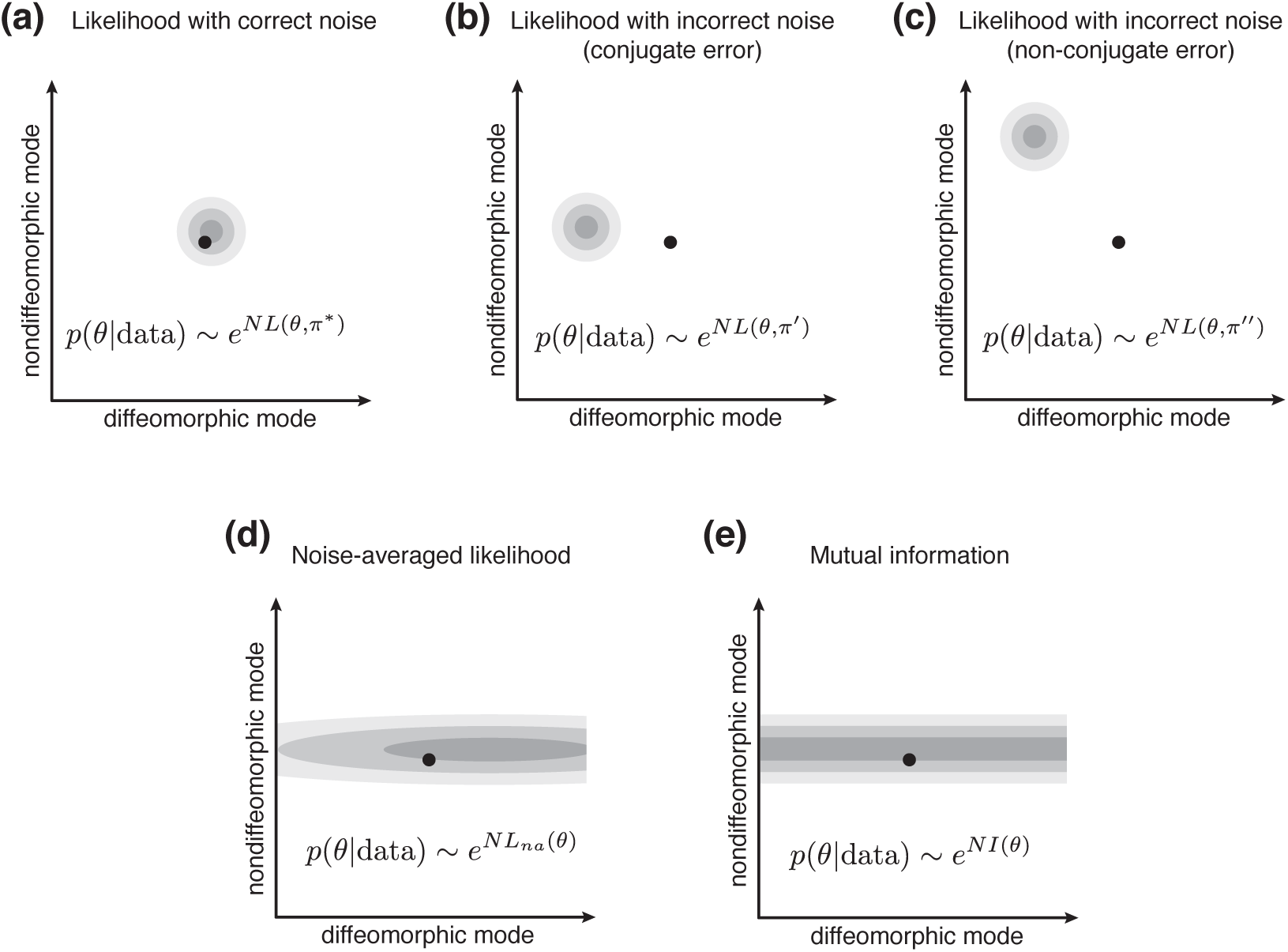
Schematic illustration of the posterior distribution *p*(*θ|* data) (gray shaded area), computed using different objective functions, in relation to the correct model *θ^*^* (dot). (a) Using likelihood with the correct noise function *π^*^* leads to inference consistent with *θ^*^*. (b) Using likelihood with a noise function *π^l^* that differs from *π^*^* only in conjugate parameters typically recovers nondiffeomorphic parameters that are consistent with *θ^*^* along with diffeomorphic parameters that are inconsistent with *θ^*^*. (c) Using likelihood with a noise function *π^ll^* that differs from *π^*^* in non-conjugate parameters typically leads to inference of *θ* that is inconsistent with *θ^*^* in both diffeomorphic and nondiffeomorphic parameters. (d) Using noise-averaged likelihood results in strong constraints on nondiffeomorphic parameters and weak constraints on diffeomorphic parameters. The constraints on the diffeomorphic parameters are determined by the choice of the noise function prior *p*(*π*). (e) Using mutual information is in principle similar to using noise-averaged likelihood, except that it provides no constraints whatsoever on diffeomorphic parameters. The problem of estimating mutual information to sufficient precision, however, remains. Figure adapted from [7].

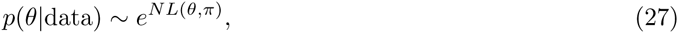

computed assuming various noise functions *π*. If one assumes the correct noise function *π^*^*, the inferred values for all parameters of *θ* will be consistent with the true parameter values of *θ^*^*. Moreover, the uncertainty in all of these parameters will decrease as *δθ ∼ N*^−1/2^.

If we instead assume an incorrect noise function, the uncertainty in all parameters will still vary as *δθ ∼ N*^−1/2^. However, the use of an incorrect noise function will lead to bias in the inferred *θ*. Which biases occur depend on which aspects of the noise function are wrong.

If we use a noise function *π^′^* that differs from *π^*^* in a parameter conjugate to a diffeomorphic parameter of *θ*, then that diffeomorphic parameter will be biased away from *θ^*^* in order to counteract the inaccuracy of *π^′^*. So likelihood-based inference will give a wrong answer, but perhaps this state of affairs isn’t too severe: the inferred model *θ^°^* will deviate from *θ^*^* in the values of parameters that we know we cannot accurately infer anyway.

However, if we use a noise function *π*′′ that differs from *π^*^* in one or more parameters that are not conjugate to any of the diffeomorphic modes of *θ*, then the inferred model *θ^o^* will, in general, deviate from *θ^*^* in the values of both diffeomorphic and nondiffeomorphic parameters. This is genuinely problematic. It means that any incorrect assumptions about the non-conjugate aspects of *π* can be expected to result in incorrect values for any of the parameters of *θ*. We therefore see that performing likelihood-based inference with an assumed noise function can, in the large *N* limit, carry a substantial risk of error propagating to the model *θ* that one wishes to learn.

The proper Bayesian approach to dealing with uncertainty in *π* is to formalize one’s uncertainty by adopting an explicit prior *p*(*π*), then computing the noise-averaged likelihood described above. This results in a situation schematized in Fig. 6d. Doing this will place tight constraints (*δθ ∼ N*^−1/2^) along all of the nondiffeomorphic modes of *θ*. However, along diffeomorphic modes one finds weak constraints that do not decrease to zero in the *N* → ∞ limit. What constraints there are arise only from the choice of the noise function prior *p*(*π*).

We therefore see that diffeomorphic model parameters and nondiffeomorphic model parameters respond to data in fundamentally different ways. Uncertainty in the values of nondiffeomorphic parameters decrease as *N* ^-1^*/*^2^ in the large *N* limit, and this is true regardless of whether one has accurate information about the noise function *π*. Uncertainty in the values of diffeomorphic parameters, however, is constrained entirely by prior knowledge of the noise function *π* when *N* is large. Unless the correct noise function is known with absolute certainty, this difference in behavior will, in the *N* →∞ limit, cause likelihood based inference to underestimate the uncertainty in all diffeomorphic parameters. This fact is a fundamental aspect of all statistical regression problems.

Unlike likelihood, which is readily computed, it is unclear how one should compute noise-averaged likelihood, or even what sort of priors *p*(*π*) might make sense. The similarity of noise-averaged likelihood and mutual information, however, suggests that one might instead perform inference using

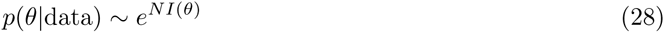

where *I*(*θ*) = *I*[*R*; *M*] is an estimate of the mutual information between model predictions *R* and measurements *M*. This was the strategy used in [5] to fit the biophysical model shown in Fig. 2. If mutual information can be accurately computed to *O*(1*/N*), then such inference will produce the constraints depicted in Fig. 6e: tight constraints on nondiffeomorphic parameters and no constraints whatsoever on diffeomorphic parameters. Doing this eliminates three problems: the need to posit an explicit noise function prior *p*(*π*), the need to integrate over all noise functions, and potential concerns about the effects of the prior *p*(*π*) on the inferred values of diffeomorphic model parameters. In this way, using mutual information in place of per-datum log noise-averaged likelihood more accurately reflects the knowledge we gain about *θ* from the data.

The downside of using Eq. (28) is that one needs to be able to estimate mutual information to an accuracy of ∼*N*^-1^. At present it remains unclear how to do this reliably. Simple approaches, such as the kernel smoothing method used in [5], do appear to work well in practice on real data. However, the accuracy of this sampling approach has yet to be systematically evaluated.

## 8 Worked example: Gaussian selection

The above principles can be illustrated in the following analytically tractable model of a Sort-Seq experiment, which we call the “Gaussian selection model”. In this model, our experiment starts with a large library of “DNA” sequences *S*, each of which is a *D*-dimensional vector drawn from a Gaussian probability distribution

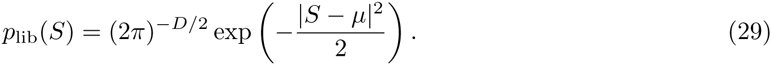

Here, *μ* is a *D*-dimensional vector defining the average sequence in the library. From this library we extract sequences into two batches, labeled *M* = 0 and *M* = 1. We fill the *M* = 0 batch with sequences sampled indiscriminately from the library, and fill the *M* = 1 batch with sequences sampled with relative probability

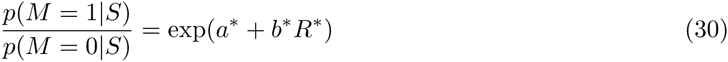

where the activity *R*^*\**^ is defined as the dot product of the sequence *S* with some “correct” model *θ^*^* (which is also a *D*-dimensional vector), i.e.

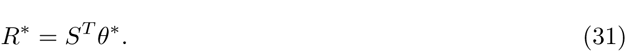

We will use *N*_*M*_ to denote the number of sequences in each batch *M*, along with *N* = *N*_0_ + *N*_1_. All of our calculations will take place in the limit where *N*_1_ is large but for which *N*_0_ is far larger. We will use *ϵ* to denote the ratio

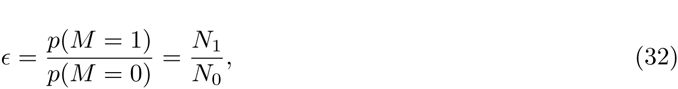

All calculations will be carried out only to first order in *ε*. This model Sort-Seq experiment is illustrated in Fig. 7.

**Fig. 7.**
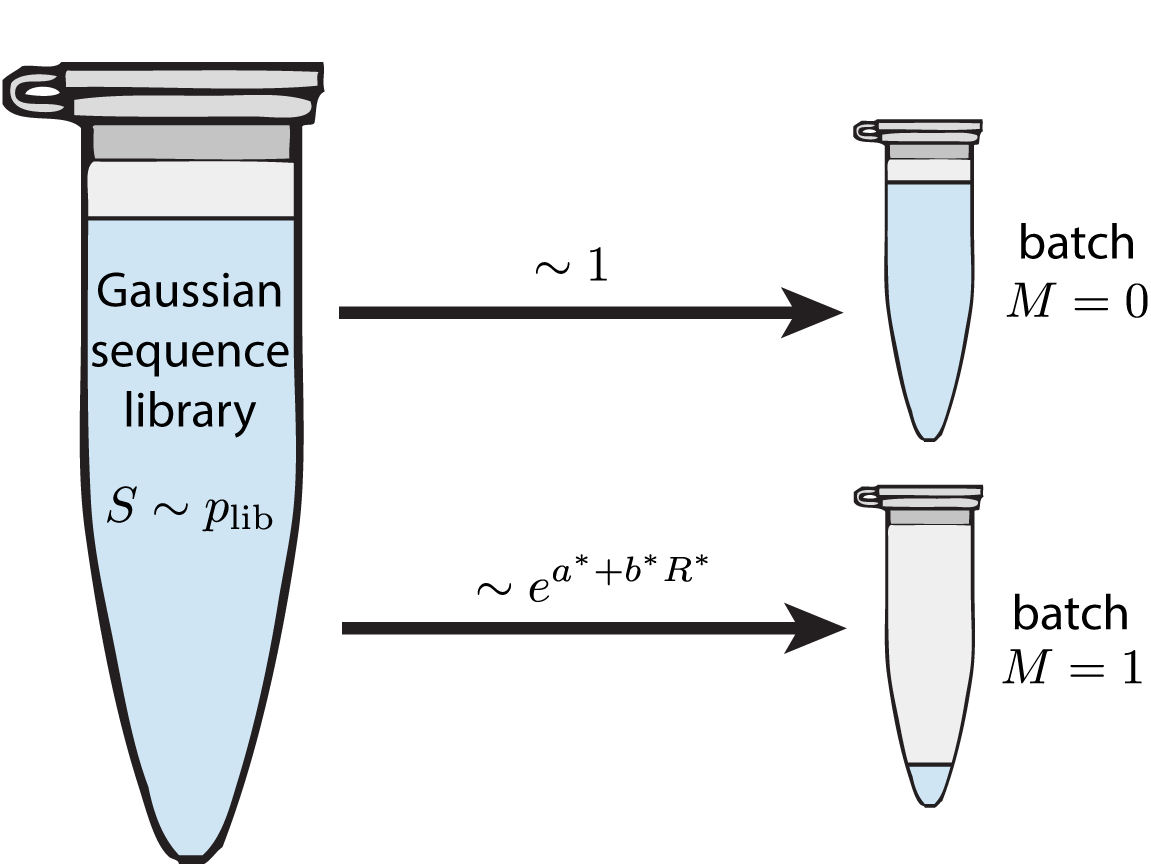
Illustration of the Gaussian selection model. We start with a library of sequences *S* distributed according to *p*_lib_(*S*). *N*_0_ sequences are sampled indiscriminately into batch 0. *N*_1_ sequences are sampled into batch 1 with relative weights exp(*a*^*\**^ + *b*^*\**^*R*^*\**^), where *R*^*\**^ = *S*^*T*^ *θ*^*\**^. All calculations are performed in the *N*_0_≫ N_*1*_limit, as suggested by the liquid volumes shown in the two tubes.

Our goal is this: given the sequences in the two batches, recover the parameters *θ^*^* defining the sequence-function relationship for *R*^*\**^. To do this, we adopt the following model for the sequence-dependent activity:

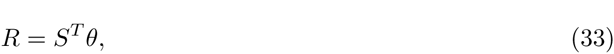

where *θ* is the *D*-dimensional vector of parameters we wish to infer, and *R* is the predicted activity. From the arguments above and in [7], it is readily seen that the magnitude of *θ*, i.e. |*θ|*, is the only diffeomorphic mode of the model: changing this parameter rescales *R*, which preserves rank order.

### 8.1 Batch-specific distributions

From this model, we can readily calculate the sequence distribution *p*(*S |M*) in each batch, as well as the model prediction distribution *p*(*R| M*) for any hypothesized model *θ*. From the selection procedure described above and illustrated in Fig. 7. Since sequences are sampled for batch 0 are indiscriminately from *P*lib,

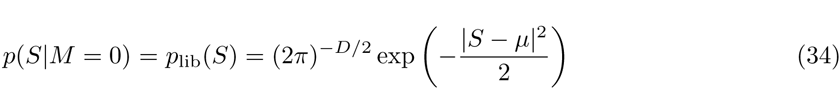

Using Bayes’s theorem, we can then use this to compute the distribution of sequences in batch 1 as follows.

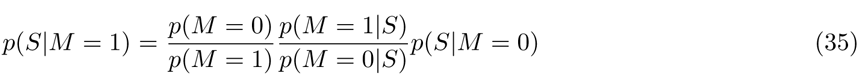

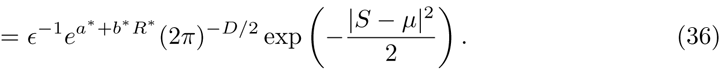

Completing the square in the exponent, we obtain

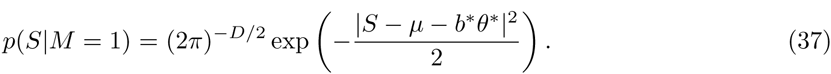

Moreover, from the normalization requirement on *p*(*S |M* = 1), we find that *c* is related to *a*^*\**^, *b*^*\**^, and *θ^*^* via

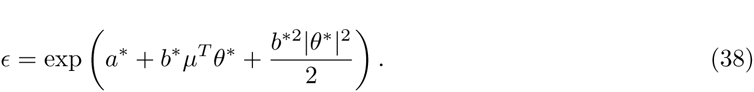

We compute the distribution of model predictions for each batch as follows. For each *M*, this distribution is defined as

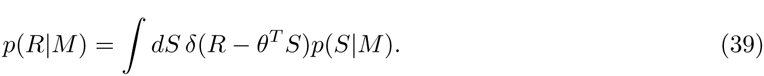

This can be analytically calculated for both of the batches owing to the Gaussian form of each. We find that

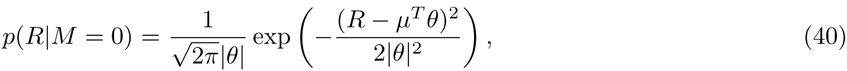

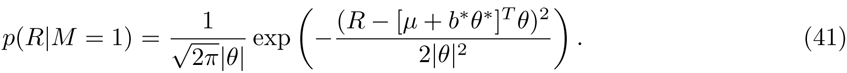

See Appendix for details.

### 8.2 Likelihood

To compute likelihood, we must posit a noise function *π*(*M |R*). Based on our prior knowledge of the selection procedure, we choose *π*(*M |R*) so that

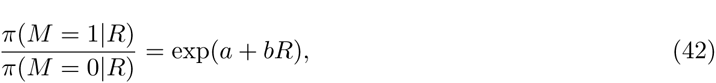

where *a* and *b* are scalar parameters that we may or may not know *a priori*. This, combined with the normalization requirement, Σ_*M*_π(*M*∣*R*) = 1 gives

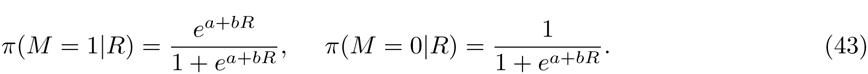

This noise function *π* is correct when *a* = *a*^*\**^ and *b* = *b*^*\**^. The parameter *b* is conjugate to the diffeomorphic mode ∣*θ*∣, while the parameter *a* is not conjugate to any diffeomorphic mode.

Using this noise function, the per-datum log likelihood *L* becomes a function of *θ*, *a*, and *b*. We compute this quantity as follows:

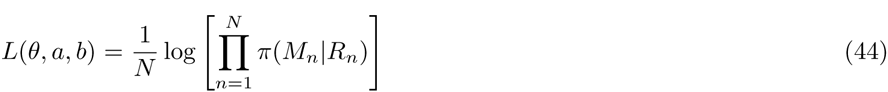

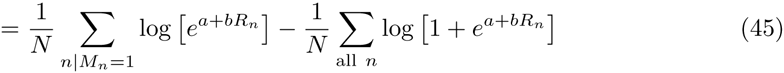

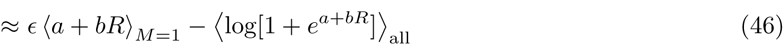

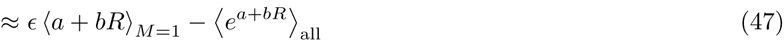

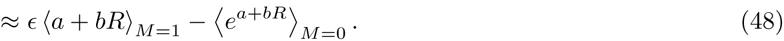

Here, *n* indexes the sequence-measurement pairs (*M*_*n*_, *S*_*n*_) that comprise our data. 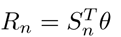 indicates the model prediction for sequence *S*_*n*_, and subscripts on the angle brackets indicate averages taken over either *p*(*S*|*M* = 1), *p*(*S*|*M* = 0), or *p*(*S*). In going from Eq. (46) to Eq. (47), we have used the assumption that *e*^*a+bR*^ ≪ 1, which we expect to hold in the *∊* → 0 limit. In the last step, we approximate the expectation over all sequences by the expectation over sequences only within batch 0. Applying the expressions for *p(R|M)* derived above, we find that the per-datum log likelihood is, in the large *N* limit, given by

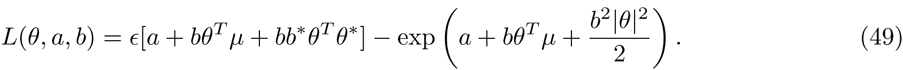

We now consider the consequences of various approaches for using *L*(*θ, a, b*) to estimate *θ^*^*. In each case, the inferred optimum will be denoted by a superscript ‘o.’ Standard regression requires that we assume a specific value for *a* and for *b*, then optimize *L*(*θ, a, b*) over *θ* alone. Setting

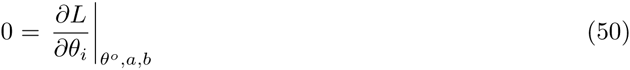

for each component *i*. By this criteria we find that the optimal model *θ^o^* is given by a linear combination of *θ^*^* and *μ*:

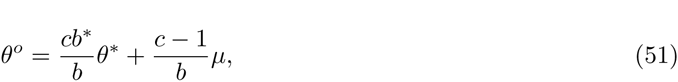

where *c* is a scalar defined by the transcendental equation

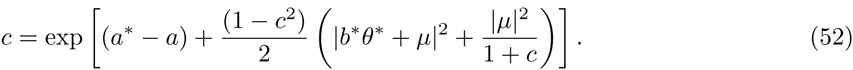

See Appendix for details. Note that *c* is determined only by the value of *a* and not by the value of *b*. Moreover, *c* = 1 if and only if *a* = *a*^*\**^.

If our assumed noise function is correct, i.e., *a* = *a*^*\**^ and *b* = *b*^*\**^, then

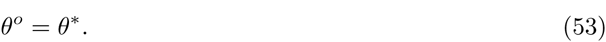

Thus, maximizing likelihood will identify the correct model parameters. This exemplifies the general behavior illustrated in Fig. 6a.

If *a* = *a*^*\**^ but *b* ≠*b* our assumed noise function will differ from the correct one only in a manner conjugate to the diffeomorphic mode *|θ|*. In this case we find that

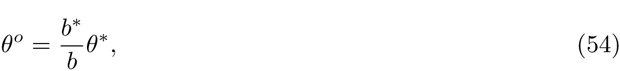

and thus *θ*^*°*^ is proportional but not equal to *θ*^*\**^. This comports with our claim above that the diffeomorphic mode of the inferred model, i.e. |*θ°*|, will be biased so as to compensate for the error in the conjugate parameter *b* of the noise function. This finding follows the behavior described in Fig. 6b.

If *a*≠*a*\*, however, *c* ≠ 1 As a result, *θ*^*°*^ is a nontrivial linear combination of *θ^*^* and *μ* and will thus point in a different direction than *θ^*^*. This is true regardless of the value of *b*. This behavior is illustrated in Fig. 6c: errors in non-conjugate parameters of the noise function will typically lead to errors in nondiffeomorphic parameters of the model.

We now consider the error bars that likelihood places on model parameters. Setting *θ* = *θ^°^* + *δθ* and expanding *L*(*θ, a, b*), we find that

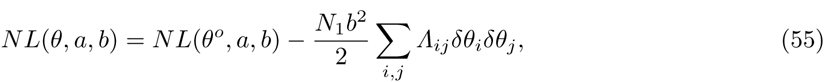

where 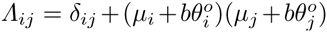. Note that all eigenvalues of *Λ* are greater or equal to 1. Adopting the posterior distribution

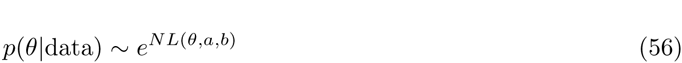

therefore gives a covariance matrix on *θ* of

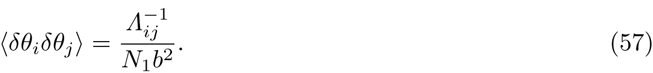

Thus, 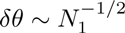 in all directions of *θ*-space. This is illustrated in Figs. 6a-c. The dependence of these fluctuations on *N*_1_ does not depend on whether the noise function *π* is correct. Therefore, when the noise function is incorrect, the finite bias introduced into *θ^°^* will cause *θ^*^* to fall outside the inferred error bars for sufficiently large *N*.

### 8.3 Mutual information

In the ϵ → 0 limit, Eq. (7) simplifies to

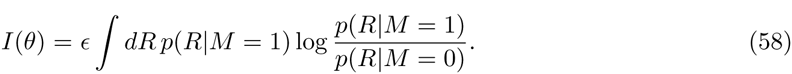

The right hand side can be evaluated exactly using Eq. (40) and Eq. (41):

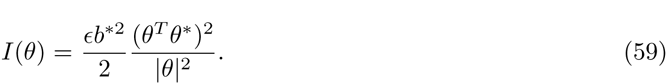

See Appendix for details. Note that the expression on the right is invariant under rescaling of *θ*. This reflects the fact that | *θ* | is a diffeomorphic mode of the model defined in Eq. (33).

To find the model *θ*^*°*^ that maximizes mutual information, we set

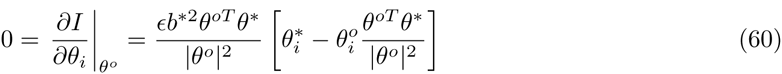

The optimal model *θ*^*°*^ must therefore be parallel to *θ^*^*, i.e.

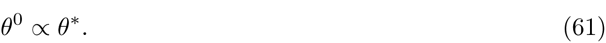

Expanding about *θ* = *θ*^*°*^ + *δθ* as above, we find that

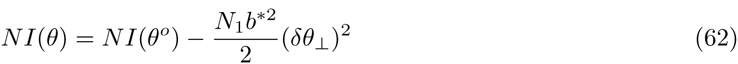

where *δθ_⊥_* is the component of *δθ* perpendicular to *θ^*^*. Therefore, if we use the posterior distribution

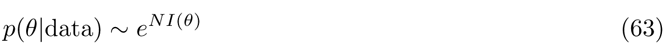

to infer *θ*, we find uncertainties in directions perpendicular to *θ^*^* of magnitude 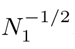. These error bars are only slightly larger than those obtained using likelihood, and have the same dependence on *N*. However, we find no constraint whatsoever on the component of *δθ* parallel to *θ^*^*. These results are illustrated by Fig. 6e

### 8.4 Noise-averaged likelihood

We can also compute the per-datum log noise-averaged likelihood, *Lna*(θ), in the case of a uniform prior on *a* and *b*, i.e. *p*(*π*) = *p*(*a, b*) = *𝒞* where *𝒞* is an infinitesimal constant. We find that

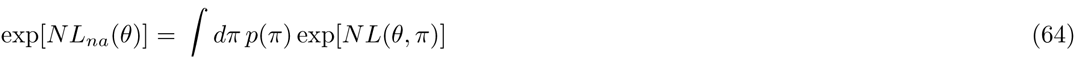

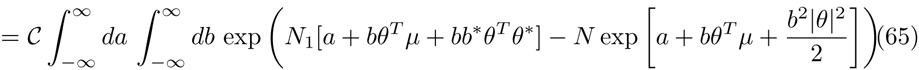

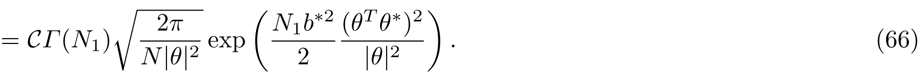

See the Appendix for details. Thus,

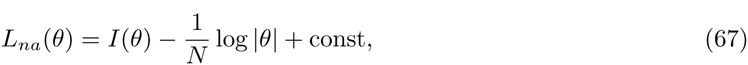

where the constant (which absorbs *𝒞* entirely) does not depend on *θ*. Therefore, if we perform Bayesian inference using noise-averaged likelihood, i.e. using

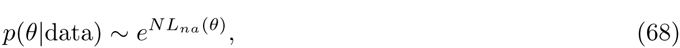

we will find in the large *N* limit that *δθ_⊥_* is constrained in the same way as if we had used mutual information, i.e.

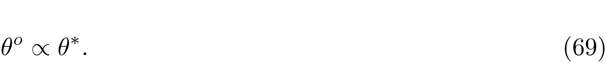

However, the noise function prior we have assumed further results in weak constraints on | *θ* | that do not tighten as *N* increases.^2^ This is illustrated schematically in Fig. 6d.

## 9 Discussion

Understanding the features of sequence-function relationships in molecular biology is going to require the ability to glean quantitative models from high-throughput experiments with poorly characterized noise. Standard regression methods require that the user assume a specific noise function up front, then find the explanatory model that has maximum likelihood. However, if the noise function one assumes is incorrect, this will in general cause errors in the sequence-function relationships that one infers.

In this paper we have shown that maximizing mutual information instead of likelihood allows one to infer sequence-function relationships from large data sets without any assumptions about the quantitative form of experimental noise. On the other hand, maximizing mutual information may fail to pin down certain model parameters. These are called diffeomorphic parameters, and are constrained by data in a fundamentally different way than are nondiffeomorphic parameters. Such nondiffeomorphic parameters can be constrained to arbitrary precision (given enough data) without any knowledge of the experimental noise function. By contrast, diffeomorphic parameters are constrained only by knowledge of the noise function and the precision of these constraints does not increase with the amount of data. The relevant aspects of inference using likelihood, mutual information, and noise-averaged likelihood, were illustrated here in an analytically tractable “Gaussian selection” model.

The development of methods for learning sequence-function relationships from Sort-Seq data and data produced by similar experiments still requires much attention. We are only in the infancy of solving this machine learning problem. One practical issue is how to accurately estimate the mutual information used for inferring model parameters, and wholly satisfactory methods for doing this have yet to be described.

1 Here, as throughout this paper, we restrict our attention to situations in which *R* is a scalar. The case of vector-valued model predictions *R* is worked out in [7].

1 In the case at hand, |*θ*^o^| is pushed all the way to zero. This is an artifact of the simple flat prior *p(a, b)*. If we instead adopt a weak Gaussian prior on b, we can still carry out the computation of *L*_*na*_ analytically, and in this case we find that |*θ*^o^| is finite.

## 10 Appendix

### 10.1 Derivation of Eqs. 40 and 41

Here we describe how to compute *p*(*R|M*) where *R* = *θ^T^ S*. We first consider the case of *M* = 0.

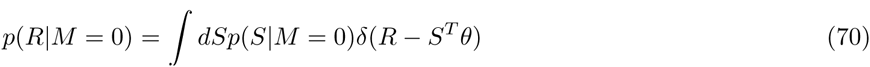

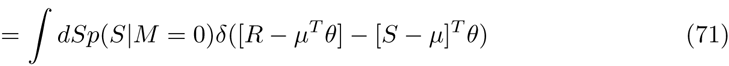

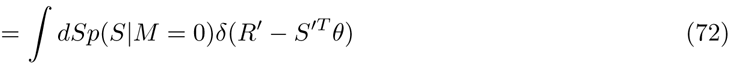

where *R*^*′*^ = *R μ^T^ θ* and *S*^*′*^ = *S μ*. Now, split *S′* up into the components parallel and perpendicular to *θ*:

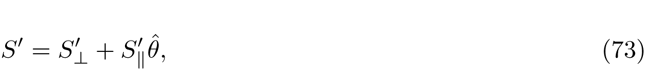

where *S*^*′*^ _*⊥*_ is a vector of dimension 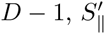 is a scalar, and 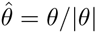 Continuing the integration,

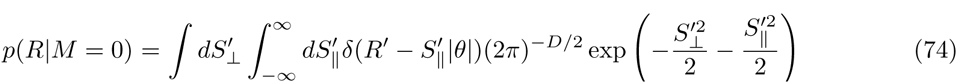

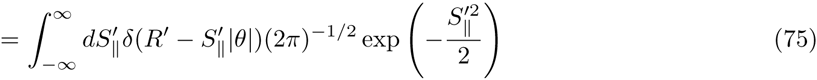

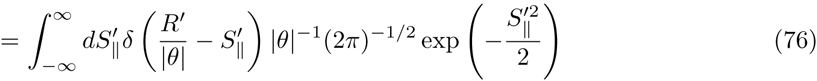

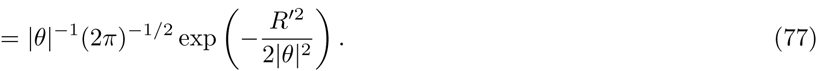

Thus we find

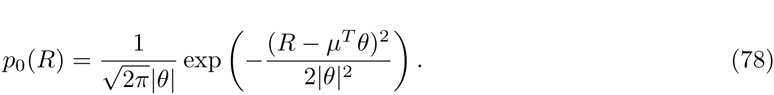

To compute *p*(*R|M* = 1), we just replace *μ → μ* + *b*^*\**^*θ*^*\**^, giving

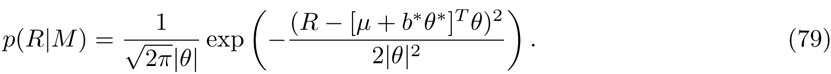

### 10.2 Derivation of Eqs. 51 and 52

Here we show how to derive the optimal *θ* for *L*(*θ, a, b*), with *a* and *b* fixed. Setting the variation of *L* to zero,

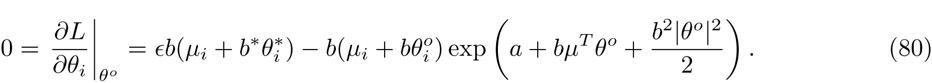

This gives

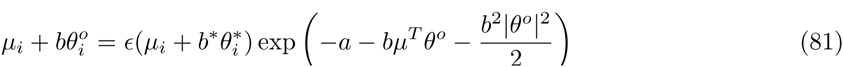

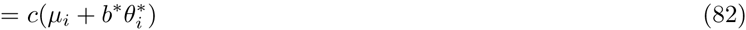

where *c* is a constant satisfying

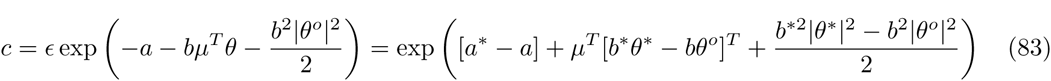

Note that the right hand side of the above equation depends implicitly on *c* through the value of *θ*.

Each component *θ_i_* must therefore satisfy

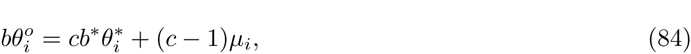

which gives Eq. (51). Using this formula for *bθ^o^*, we find that

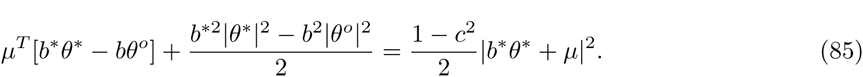

Plugging this into Eq. (83) gives Eq. (52).

### 10.3 Derivation of Eqs. 58 and 59

We derive Eq. (58) as follows. To ease notation a bit, we define *p*_*M*_ (*R*) = *p*(*R|M*).

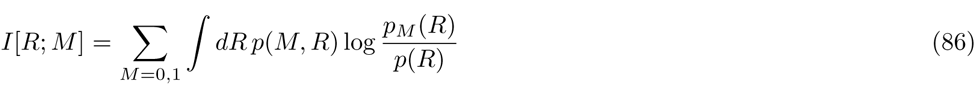

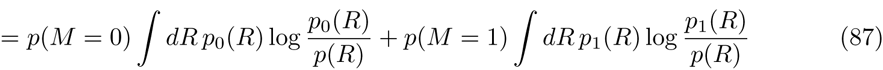

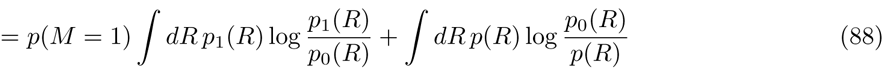

the first term in Eq. (88) is the right hand side of Eq. (58). We will now show that the second term is of order ϵ^2^, and can therefore be ignored. Rearranging

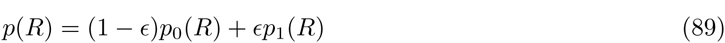

gives

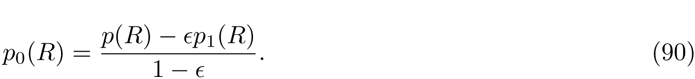

Plugging this into the second term of Eq. (88) gives

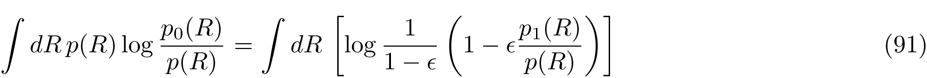

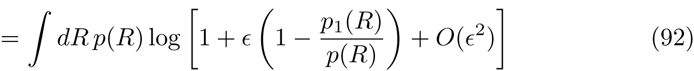

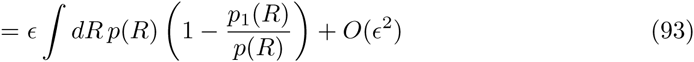

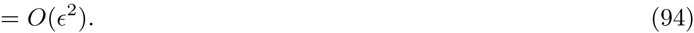

Eq. (59) is derived as follows:

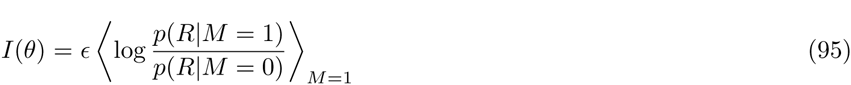

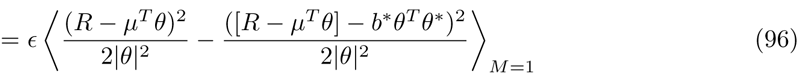

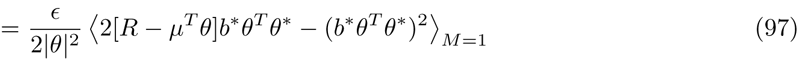

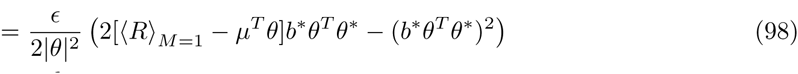

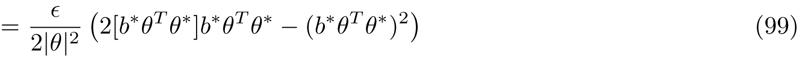

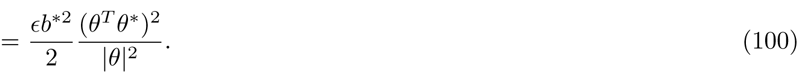

### 10.4 Derivation of Eq. 66

Here we show how to evaluate the equation, Eq. (65), for the noise-averaged likelihood *e*^*NL*_*na*_^(*θ*). First, interchange the order of integration and define *a*^*’*^ = *a* + *bθ^T^ μ*. This gives,

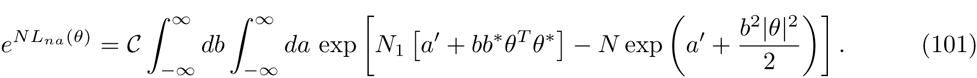

Next, define 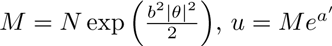, and 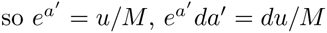. This gives

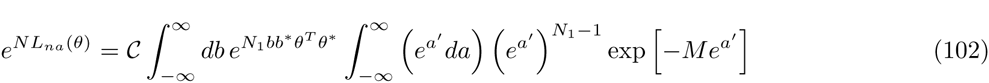

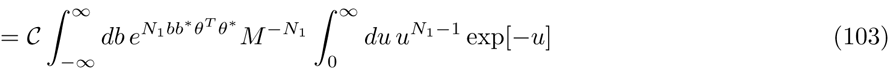

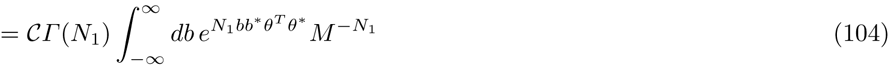

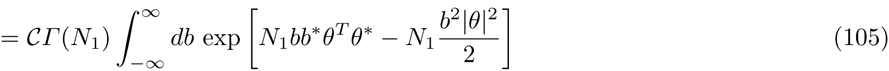

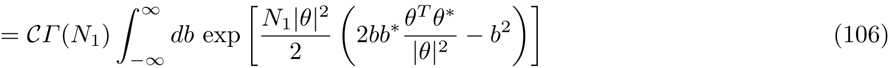

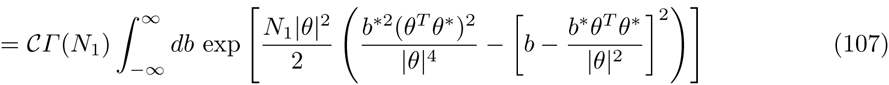

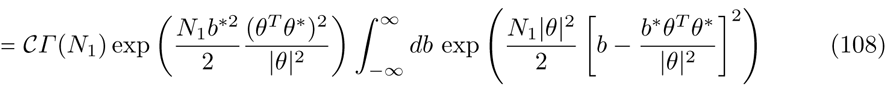

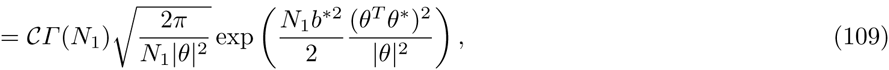

which is Eq. (66).

## Acknowledgements

We would like to thank L. Peliti, O. Revoire, and T. Mora for organizing this special issue. This work was supported by the Simons Center for Quantitative Biology at Cold Spring Harbor Laboratory.

